# Prenatal DEHP plastic chemical exposure increases the likelihood of child autism and ADHD symptoms through epigenetic programming

**DOI:** 10.1101/2025.03.03.641339

**Authors:** Samuel Tanner, Alex Eisner, Boris Novakovic, Lada Holland, Toby Mansell, Gillian England-Mason, Sarah Merrill, Deborah Dewey, Martin O’Hely, Christos Symeonides, Richard Saffery, Jochen Mueller, Mimi LK Tang, Peter D Sly, Peter Vuillermin, the BIS Investigator Group, Chol-Hee Jung, Daniel Park, Anne-Louise Ponsonby

## Abstract

Increasing evidence implicates prenatal exposure to di-(2-ethylhexyl) phthalate (DEHP), a common endocrine-disrupting plastic chemical, in autism spectrum disorder (ASD) and attention-deficit/hyperactivity disorder (ADHD). However, the underlying mechanisms are poorly understood. Here we examined whether cord blood DNA methylation, a key epigenetic marker, mediates the association between prenatal DEHP exposure and ASD/ADHD symptoms in 847 children enrolled in the Barwon Infant Study. ASD and ADHD are complex phenotypes characterised by differences at the gene regulatory network and neuronal circuit level, where heterogeneous genetic and environmental risk factors converge. Accordingly, we employed a data-driven computational strategy that helped elucidate broader functional epigenetic signatures of ASD and ADHD elicited by DEHP exposure. This included (1) a methylation profile score for DEHP exposure (MPS_DEHP_), and (2) an analysis of co-methylated gene networks. Causal mediation analysis demonstrated that both MPS_DEHP_ and a DEHP-associated network of co-methylated genes mediated the effect of DEHP exposure on increased ASD and ADHD symptoms at ages 2 and 4 years (proportion of effect mediated ranged from 0.21 to 0.80). The co-methylation network was enriched for neural cell-type markers, ASD risk genes (including *FOXP1*, *SHANK2,* and *PLXNB1*), and targets of endocrine receptors previously linked to DEHP (including targets of the estrogen receptor ERα and the glucocorticoid receptor GR), providing biological plausibility. We validated key results in independent blood (n=66) and postmortem brain (n=40) DNA methylation datasets. These findings provide mechanistic evidence linking DEHP to ASD and ADHD symptoms and reinforce growing concerns regarding the risks of prenatal exposure.

**Significance:** Exposure to endocrine-disrupting plastic chemicals has been linked to adverse neurodevelopment, but the underlying biological mechanisms remain unclear. We demonstrate that prenatal exposure to di-(2-ethylhexyl) phthalate (DEHP), a common plasticizer, increases autism and ADHD symptoms through alterations in DNA methylation, a key epigenetic regulator of gene activity. Using birth cohort data, we identify epigenetic signatures of prenatal DEHP exposure, including alterations in an endocrine-related co-methylation network enriched for neural cell-type markers and known autism-associated genes. These signatures mediate the effects of DEHP on autism and ADHD symptoms and are also associated with autism in external blood and postmortem-brain datasets, providing independent validation. This causal evidence further underscores concerns regarding the consequences of prenatal plastic-chemical exposure on the developing brain.

## 1 Introduction

Autism spectrum disorder (ASD) and attention-deficit/hyperactivity disorder (ADHD) are related and often co-occurring neurodevelopmental conditions with an estimated global prevalence of at least 1-2% and 2-3%, respectively.^1,2^ ASD is characterised by restricted or repetitive behaviours and challenges in social communication, and ADHD by inattention, impulsivity, and/or hyperactivity.^3^ The aetiologies of ASD and ADHD are multifactorial, reflecting a complex interplay of genetic risk and early environment. Their prevalence appears to be increasing, even after accounting for improved awareness and changes in diagnostic criteria.^3^ The rate of increase is too rapid to be explained by genetics alone.^4^ Consequently, concern is mounting around the role of environmental exposures identified in epidemiological studies as risk factors.

Phthalate plasticisers are emerging as an important class of environmental risk factors. Phthalates are added to common household plastics to improve flexibility but are not covalently bonded to the polymer and readily leach out.^5^ Of particular concern is di-(2-ethylhexyl) phthalate (DEHP), an endocrine disruptor that can interfere with hormonal pathways critical to early brain development, including estrogen and androgen signalling.^6^ DEHP has been linked to adverse neurodevelopment, including ASD and ADHD symptoms, in multiple human and controlled animal studies,^7–10^ but its mode of action remains unclear and causal evidence from human studies is minimal.

Several lines of evidence indicate that DEHP may impact neurodevelopment through epigenetic mechanisms. Epigenetics, literally meaning “above genetics”, encompasses a range of mechanisms that facilitate changes in gene expression without modifying the underlying DNA sequence.^11^ Importantly, epigenetic processes are dynamic and sensitive to the environment, providing an interface through which environmental factors can influence intrinsic genetic programmes.^12^ The best-characterised epigenetic layer in humans is DNA methylation, involving the addition of a methyl group to a cytosine-guanine dinucleotide (CpG site) on the DNA molecule. DNA methylation has diverse regulatory functions, such as silencing target genes by blocking the binding of transcription factors. It plays fundamental roles in the transcriptional programming of the developing brain, modulating gene expression spatially and temporally in response to genetic and environmental cues.^13^

DNA methylation also represents a point of susceptibility to adverse exposures. We have previously shown that in males, prenatal exposure to another endocrine-disrupting plastic chemical, bisphenol A (BPA), may increase autism symptoms in part through hypermethylation of the aromatase-gene brain promoter.^14^ Epigenome-wide association studies for prenatal DEHP exposure also reveal alterations in DNA methylation, including around neuronal genes. ^15^ Similarly, differences in DNA methylation have been identified in individuals with ASD and ADHD in both peripheral and postmortem-brain tissue.^16–18^ However, this evidence is yet to be synergised through an examination of DNA methylation more specifically as a causal mediator between DEHP exposure and ASD or ADHD symptoms. Advances in causal inference methodologies now make this feasible even in observational studies provided that DEHP exposure, DNA methylation, and neurodevelopmental outcomes are measured in the same individuals.^19^

Crucially, growing evidence points to ASD and ADHD as systems-level disorders, with a constellation of genetic and environmental risk factors converging on shared gene networks and neuroanatomical structures.^20,21^ It is the perturbation of these systems-level features that underlies ASD and ADHD symptomologies. Similarly, environmental toxins commonly disrupt DNA methylation over functional gene networks rather than at CpG sites in isolation or stochastically across the genome.^22^ Systems biology approaches are therefore required to characterise higher-level functional epigenetic signatures of ASD and ADHD altered in response to DEHP exposure that might be missed by traditional CpG- or gene-level analyses.

Methylation profile scores (MPSs) and co-methylation network (Co-MN) analysis are two powerful systems-biology approaches that can help elucidate such signatures. The epigenetic analogue of polygenic risk scores, MPSs summarise methylation across multiple CpG sites associated with a condition of interest, resulting in a single score for each participant.^23^ MPSs can also encode signatures of environmental exposures, such as DEHP, and have been employed previously to examine causal mediation effects distributed across the epigenome.^24^ Co-methylation network analysis leverages the correlation structures in DNA methylation datasets to identify clusters of co-regulated CpG sites.^25^ The rationale for this approach is that DNA methylation is generally not altered at CpG sites in isolation but across CpG networks as genes within shared pathways are activated or inhibited together. Co-methylation network analysis can yield rich functional insights.^25^

Here, using data from the Barwon Infant Study, a population-derived birth cohort, we assess cord-blood DNA methylation as a causal mediator between higher prenatal DEHP exposure and ASD and ADHD symptoms (ages 2 and 4 years) at two levels: (1) through a methylation profile score for DEHP exposure, and (2) through co-methylation networks identified using Weighted Gene Correlation Network Analysis (WGCNA). We strengthen the argument for causation by triangulating across multiple lines of evidence,^26^ highlighting the likely mechanisms involved and replicating key findings in independent blood and postmortem-brain DNA methylation datasets.

## 2 Results

### 2.1 Study characteristics and analysis workflow

The Barwon Infant Study (BIS) is an extensively profiled birth cohort of 1074 children. Study characteristics and the analysis workflow are outlined in Figure 1. The BIS cohort is described further in Supplementary Table 1 and Figure 1. Prenatal DEHP exposure measures (at 36 weeks of gestation) were available for 847 BIS participants, of whom over 98% had detectable levels of DEHP.^27^

**Fig. 1.**
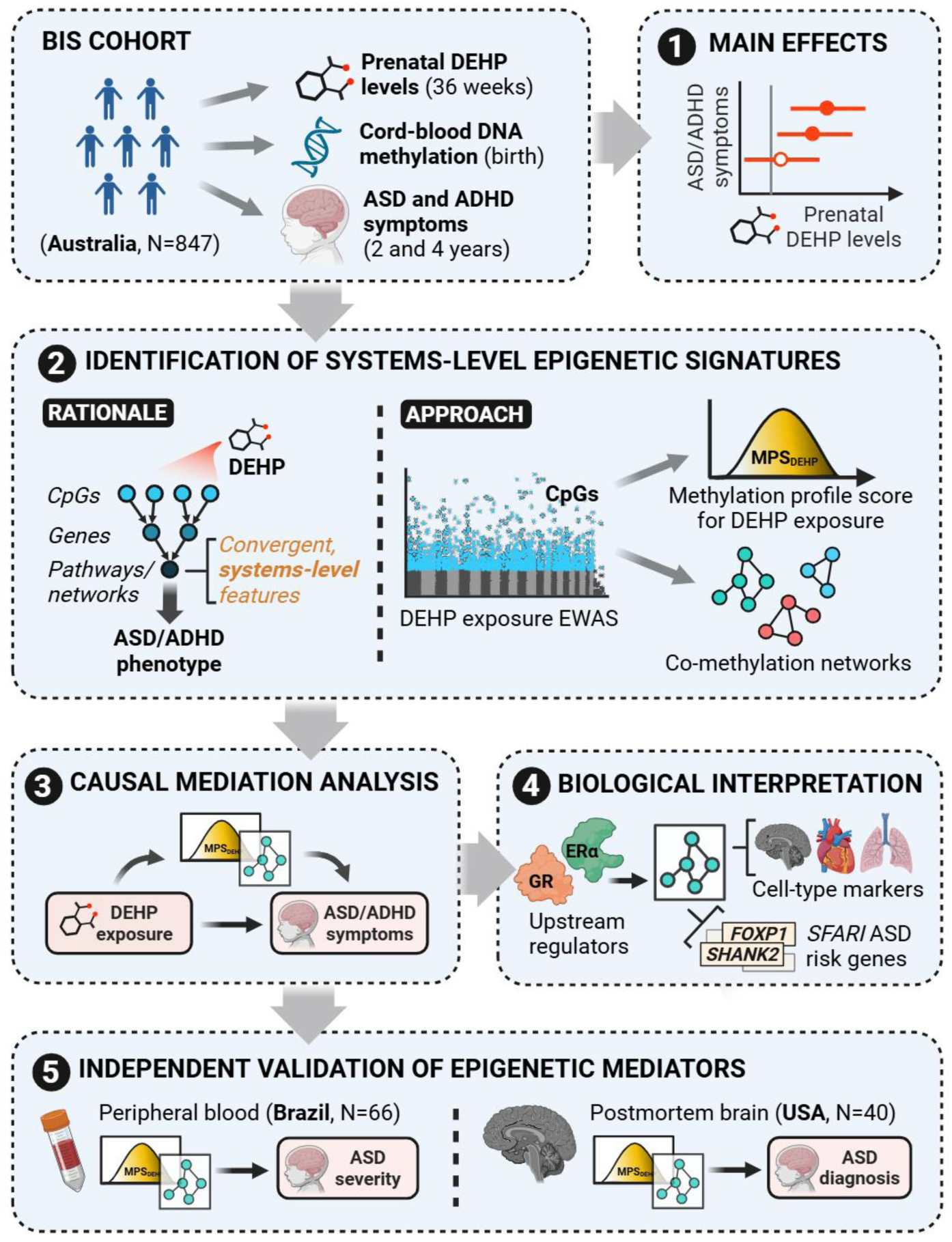
Schematic representation of the workflow for identifying epigenetic alterations mediating the association between prenatal DEHP exposure and ASD/ADHD symptoms. The workflow consisted of five key steps: (1) assessment of *main effects* of prenatal DEHP levels on neurodevelopmental outcomes; (2) identification of *systems-level epigenetic signatures* by aggregating results from an epigenome-wide association study (EWAS) for DEHP exposure using a methylation profile score (MPS) and an investigation of co-methylated gene networks; (3) *causal mediation analysis* to test the mediating role of the identified epigenetic signatures in DEHP-ASD/ADHD associations; (4) *biological interpretation* through enrichment analyses of co-methylated gene networks, including identification of upstream regulators, cell-type markers, and ASD risk genes; and (5*) independent validation of epigenetic mediators* in peripheral blood and postmortem brain DNA methylation datasets.

ASD and ADHD symptoms were assessed at age 2 years on 676 participants using the Child Behavior Checklist for ages 1.5–5 (CBCL_2yrs_)^28^ and at age 4 years on 791 participants using the Strengths and Difficulties Questionnaire P4-10 (SDQ_4yrs_),^29^ both previously validated for this purpose.

### 2.2 Prenatal DEHP exposure is associated with increased ASD and ADHD symptoms

We first examined the association between continuous prenatal DEHP exposure levels and the ASD- and ADHD-related neurodevelopmental outcomes. With adjustment for potential confounders (see Figure 2 caption), prenatal DEHP exposure was associated with increased SDQ_4yrs_ peer problems (*β* = 0.142 per doubling in the daily maternal intake of DEHP [95% CI 0.033, 0.251], *p* = 0.011), with weaker but consistent evidence for other symptoms scales (Figure 2a).

**Fig. 2.**
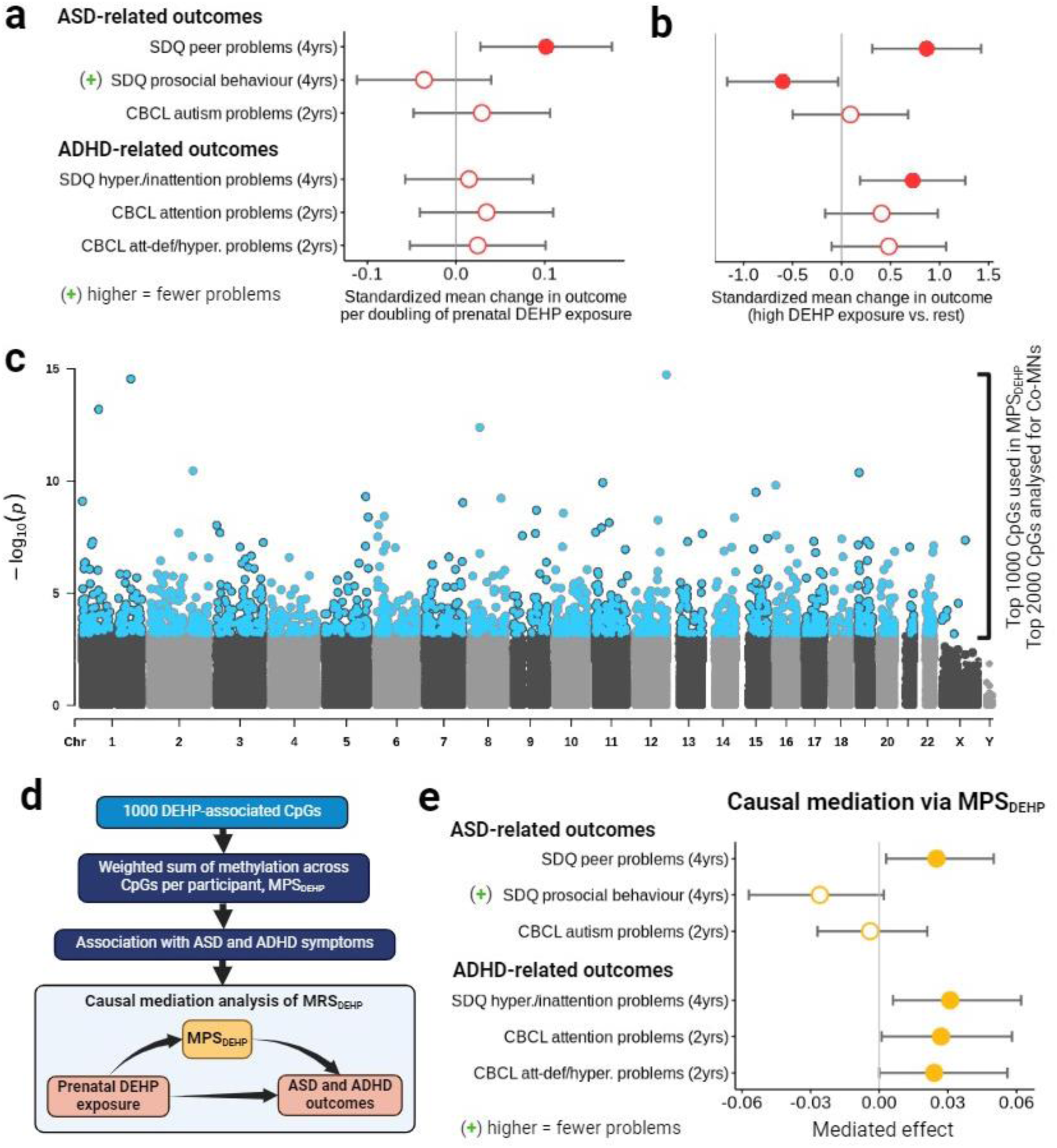
DEHP’s main effects on neurodevelopmental outcomes and its mediation via a methylation profile score. **a, b** Association between prenatal DEHP exposure levels (continuous measure and high vs. rest (DEHP_HIGH_), respectively) and ASD- and ADHD-related outcomes at ages 2 and 4 years, adjusting for age, sex, genetic ancestry, multiparity, maternal education, and household income (details in Methods). **c** Manhattan plot showing an epigenome-wide association study to screen for CpG sites associated with DEHP_HIGH_. Rather than assessing each CpG site individually as a mediator of DEHP’s effects, we aggregated high-ranking CpG sites using two systems-biology approaches: (i) a methylation profile score for DEHP exposure (MPS_DEHP_) (using the top 1000 DEHP_HIGH_-associated CpGs) and (ii) a co-methylation network (Co-MN) analysis (using the top 2000 CpGs). We then analysed these systems-level factors for mediated effects. **d** Overview of mediation analysis using MPS_DEHP_. **e** Effect of prenatal DEHP exposure (continuous measure) on ASD- and ADHD-related symptoms (scaled for plotting) transmitted via MPS_DEHP_, estimated using causal mediation analysis.^33^ Mediation models were adjusted for age, sex, genetic ancestry, multiparity, seven cord blood cell-type proportions, maternal education, and household income (details in Methods). Cord-blood samples with maternal contamination were excluded from the analysis.

In previous work, we have reported that clearer positive associations were evident between DEHP and these outcomes when children were classified with a threshold indicating high exposures.^30^ Here, to examine high-dose effects, we defined a secondary high-DEHP exposure variable (DEHP_HIGH_) representing the top 2% of exposure vs. the rest. DEHP_HIGH_ demonstrated markedly stronger associations with ASD and ADHD symptoms, including increased SDQ_4yrs_ peer problems (*β* = 1.416 for high exposure vs. rest [95% CI 0.602, 2.231], *p* = 0.0007), reduced SDQ_4yrs_ prosocial behaviour (*β* = −1.123 [95% CI −2.156, −0.091], *p* = 0.033), and increased SDQ_4yrs_ hyperactivity/inattention problems (*β* = 1.690 [95% CI 0.452, 2.928], *p* = 0.008) (Figure 2b).

Given that causal mediation analysis has greater power than main-effect models to detect underlying exposure-outcome relationships,^31^ all six outcomes were examined using causal mediation analysis.

### 2.3 Epigenome-wide association study for prenatal DEHP exposure identifies CpG sites for systems-level mediation analyses

To investigate whether DNA methylation mediates the neurotoxic effects of DEHP exposure, we analysed epigenome-wide methylation data profiled in cord blood at birth using the Illumina EPIC 850k array. After pre-processing, the methylation status for 798,259 CpG sites was available for 868 participants, 695 of which also had prenatal DEHP exposure measured. We screened for DEHP-associated CpG sites by conducting an epigenome-wide association study (EWAS) for prenatal DEHP exposure (Figure 2c). While we used the continuous DEHP exposure measure for all subsequent analyses, we used DEHP_HIGH_ for this screening EWAS to improve power to detect effects at the individual-CpG level.^32^ We ranked CpG sites by statistical significance and analysed high-ranking CpG sites for systems-level mediators using the two approaches described below. A simulation analysis demonstrated that this strategy of constructing composite mediators from individual exposure-associated CpG sites does not bias mediation results towards significance (Supplementary Figures 2 and 3).

### 2.4 Epigenome-wide alterations in DNA methylation mediate the association between prenatal DEHP exposure and increased ASD and ADHD symptoms

#### 2.4.1 Methylation profile score for prenatal DEHP exposure

To estimate whether DNA methylation differences at birth distributed across the epigenome mediate the association between prenatal DEHP exposure and childhood ASD and ADHD symptoms, we constructed a methylation profile score for prenatal DEHP exposure (MPS_DEHP_) using the top 1,000 CpGs from the DEHP_HIGH_ EWAS (Figure 2d). MPS_DEHP_ was calculated for each participant by summing each CpG’s methylation level weighted by its estimated effect size for DEHP_HIGH_, meaning that higher-confidence CpGs with stronger effect estimates contribute most to the score. MPS_DEHP_ was positively associated with the continuous prenatal DEHP exposure variable (*β* = 0.33, *p* = < 2×10^-16^).

#### 2.4.2 Association between methylation profile score and ASD and ADHD symptoms

We next examined the association between MPS_DEHP_ and ASD and ADHD symptoms to determine whether MPS_DEHP_ is a potential mediator. MPS_DEHP_ was associated with subsequent increased SDQ_4yrs_ peer problems (*β* = 0.13, *p* = 0.006), reduced SDQ_4yrs_ prosocial behaviour (*β* = -0.10, *p* = 0.008), and increased SDQ_4yrs_ hyperactivity/inattention problems (*β* = 0.12, *p* = 0.014) and CBCL_2yrs_ attention problems (*β* = 0.09, *p* = 0.038).

#### 2.4.3 Causal mediation analysis of methylation profile score

Since MPS_DEHP_ was associated with both prenatal DEHP exposure and adverse neurodevelopment, we applied causal mediation analysis to partition the total effect of prenatal DEHP exposure (continuous measure) on each of the ASD and ADHD symptoms scales (shown in Figure 2a, b) into an estimated *indirect effect* (mediated via MPS_DEHP_) and *direct effect* (operating through other biological mechanisms). After adjustment, MPS_DEHP_ was shown to mediate the effect of DEHP exposure on increased SDQ_4yrs_ peer problems (*β_ACME_* = 0.025 [95% CI 0.003, 0.050], *p* = 0.020, proportion mediated = 0.25), SDQ_4yrs_ hyperactivity/inattention problems (*β_ACME_* = 0.031 [95% CI 0.006, 0.062], *p* = 0.016, proportion mediated undefined (see Methods)), CBCL_2yrs_ attention problems (*β_ACME_* = 0.027 [95% CI 0.001, 0.058], *p* = 0.040, proportion mediated = 0.80), and CBCL_2yrs_ attention-deficit/hyperactivity problems (*β_ACME_* = 0.024 [95% CI 0.0001, 0.056], *p* = 0.048, proportion mediated undefined (see Methods)) (Figure 2e). These findings suggest that DEHP increases ASD and ADHD symptoms in part by eliciting coordinated, genome-wide alterations in DNA methylation.

### 2.5 DNA co-methylation network mediates the association between prenatal DEHP exposure and increased ASD and ADHD symptoms

#### 2.5.1 Identification of DEHP-related co-methylation networks

To gain more mechanistic insights into DEHP’s mode of action, we next investigated whether CpG Co-MNs mediate DEHP’s effects on ASD and ADHD symptoms. To this end, we analysed the top 2000 DEHP-associated CpGs using WGCNA, a data-driven, correlation-based algorithm for biological network construction.^34^ We analysed 2000 rather than 1000 CpGs because, unlike MPS_DEHP_, WGCNA filters out CpG sites that do not fall into correlated networks, thus helping to eliminate false positives. WGCNA detected 10 networks of co-methylated CpGs (Figure 3a, b), ranging in size from 36 to 791 CpGs. Each Co-MN was assigned a numeric label and a colour for convenience and represented for statistical analysis by its first principal component (Figure 3c).

**Fig. 3.**
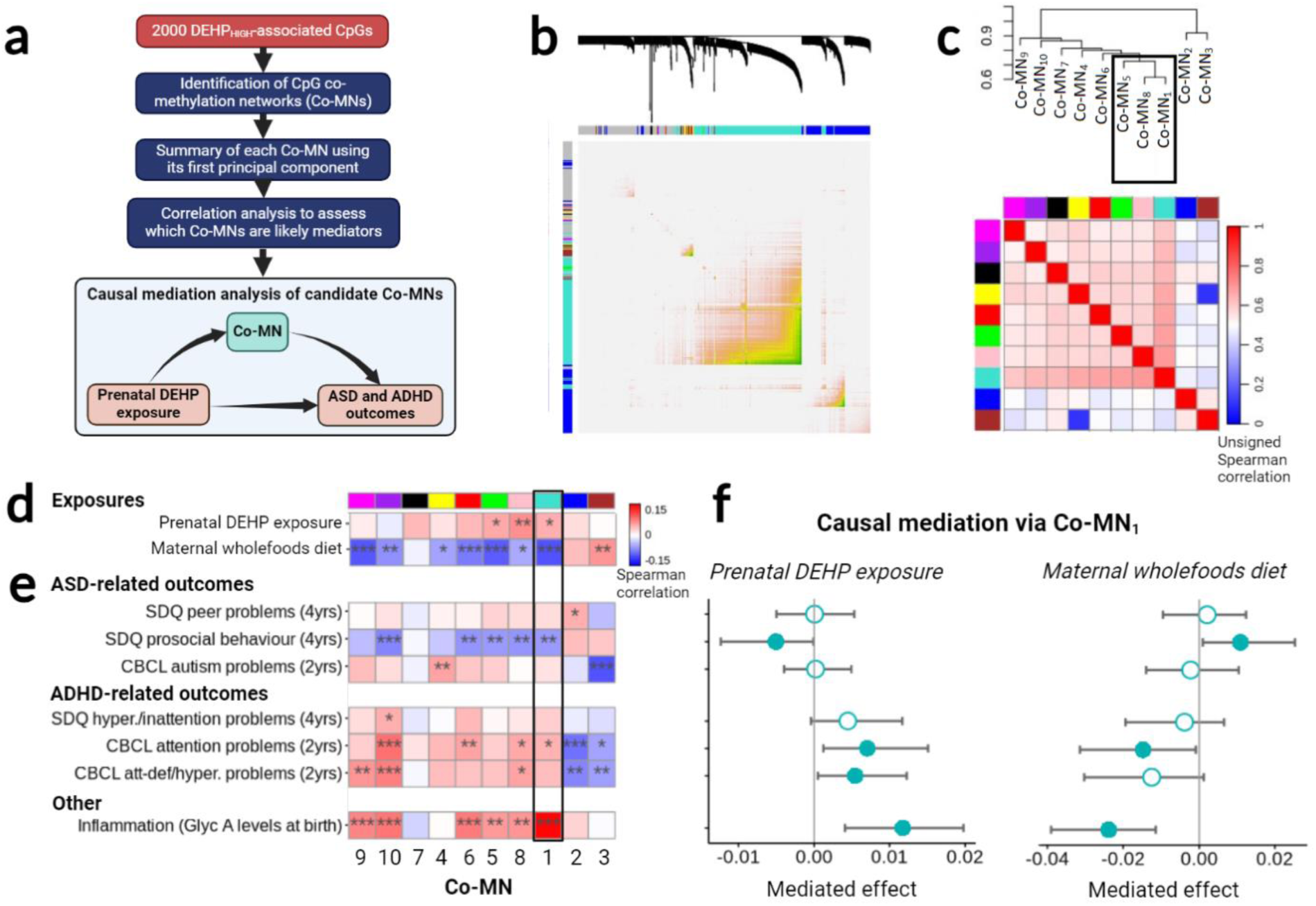
Investigation of co-methylated CpG networks mediating the effects of prenatal DEHP exposure. **a** *Summary of the procedure used to identify co-methylation networks (Co-MNs) among the 2000 DEHP-associated CpG sites*. Ten Co-MNs were identified. Each Co-MN was assigned a unique numeric label (ordered by Co-MN size) and a colour for convenience and represented for causal mediation analysis using its first principal component. **b** Heatmap of correlations among the 2000 DEHP-associated CpG sites, demonstrating the strong intercorrelations within each network. **c** Clustering of the Co-MNs themselves (represented by their first principal components), showing their higher-level relationships. **d, e** Correlation analysis of the ten Co-MNs (with their order corresponding to the clustering in) **c**) against the exposures and ASD and ADHD symptoms outcomes of interest to screen for potential mediators. The most likely mediator emerged as Co-MN_1_ (turquoise), consisting of 791 CpG sites and correlated with both prenatal DEHP exposure and two neurodevelopmental outcomes. **f** Effect transmitted via Co-MN_1_ of (left) prenatal DEHP exposure and (right) a maternal wholefoods diet on the ASD- and ADHD-related outcomes (scaled for plotting), estimated using causal mediation analysis. With DEHP as exposure, mediation models were adjusted for age, sex, genetic ancestry, multiparity, seven cord blood cell-type proportions, maternal education, and household income (details in Methods). Additional adjustment for wholefoods diet did not alter these findings. With wholefoods diet as exposure, the same covariate set was used except that granulocyte and nucleated red blood cell-type proportions were omitted from the mediation models as they partly mediated the effect of diet on Co-MN_1_.

#### 2.5.2 Screening of co-methylation networks for potential mediators

We defined potential mediators as Co-MNs that correlate with both prenatal DEHP exposure (continuous measure) and at least one neurodevelopmental outcome (Figure 3d, e). Co-MN_1_ (791 CpGs), Co-MN_8_ (78 CpGs), and Co-MN_5_ (36 CpGs) met these criteria. However, correlations for Co-MN_8_ and Co-MN_5_ were strongly influenced by outliers.

We thus focused on Co-MN_1_, which was also the largest network. Adjusted linear regression confirmed the association between Co-MN_1_ and both prenatal DEHP exposure (*β* = 0.003, *p* = 0.002) and adverse neurodevelopment, including reduced SDQ_4yrs_ prosocial behaviour (*β* = 2.60, *p* = 0.027) and increased CBCL_2yrs_ attention problems (*β* = 3.09, *p* = 0.014).

#### 2.5.3 Causal mediation analysis of candidate co-methylation network

We then tested Co-MN_1_ using causal mediation analysis. Co-MN_1_ mediated the effect of prenatal DEHP exposure (continuous measure) on reduced SDQ_4yrs_ prosocial behaviour (*β_ACME_* = −0.008 [95% CI −0.022, −0.00003], *p* = 0.048, proportion mediated = 0.21), and increased CBCL_2yrs_ attention problems (*β_ACME_* = 0.011 [95% CI 0.002, 0.024], *p* = 0.008, proportion mediated = 0.21) and CBCL_2yrs_ attention-deficit/hyperactivity problems (*β_ACME_* = 0.013 [95% CI 0.002, 0.029], *p* = 0.024, proportion mediated = 0.21), with consistently adverse effect directions for other outcomes (Figure 3f). Thus, using an alternative approach tied more closely to biological function, we also identified a systems-level epigenetic signature mediating the adverse effect of DEHP on early child neurodevelopmental outcomes.

### 2.6 Co-methylation network Co-MN_1_, mediating DEHP’s effects, is enriched for hormone-receptor targets, neural cell-type markers, and ASD risk genes

To shed light on the function of Co-MN_1_, we mapped each of the 791 CpGs in the network to its nearest gene and performed functional enrichment analyses on the resulting set of 531 genes. The network’s most central (i.e. intercorrelated) CpG site, cg19126690, was located in *NFIA* (Figure 4a), a known ASD and ADHD risk gene.^35^ *NFIA* encodes a transcription factor critical in early neurodevelopment, including in the formation of the corpus callosum and cerebellum.^36^

**Fig. 4.**
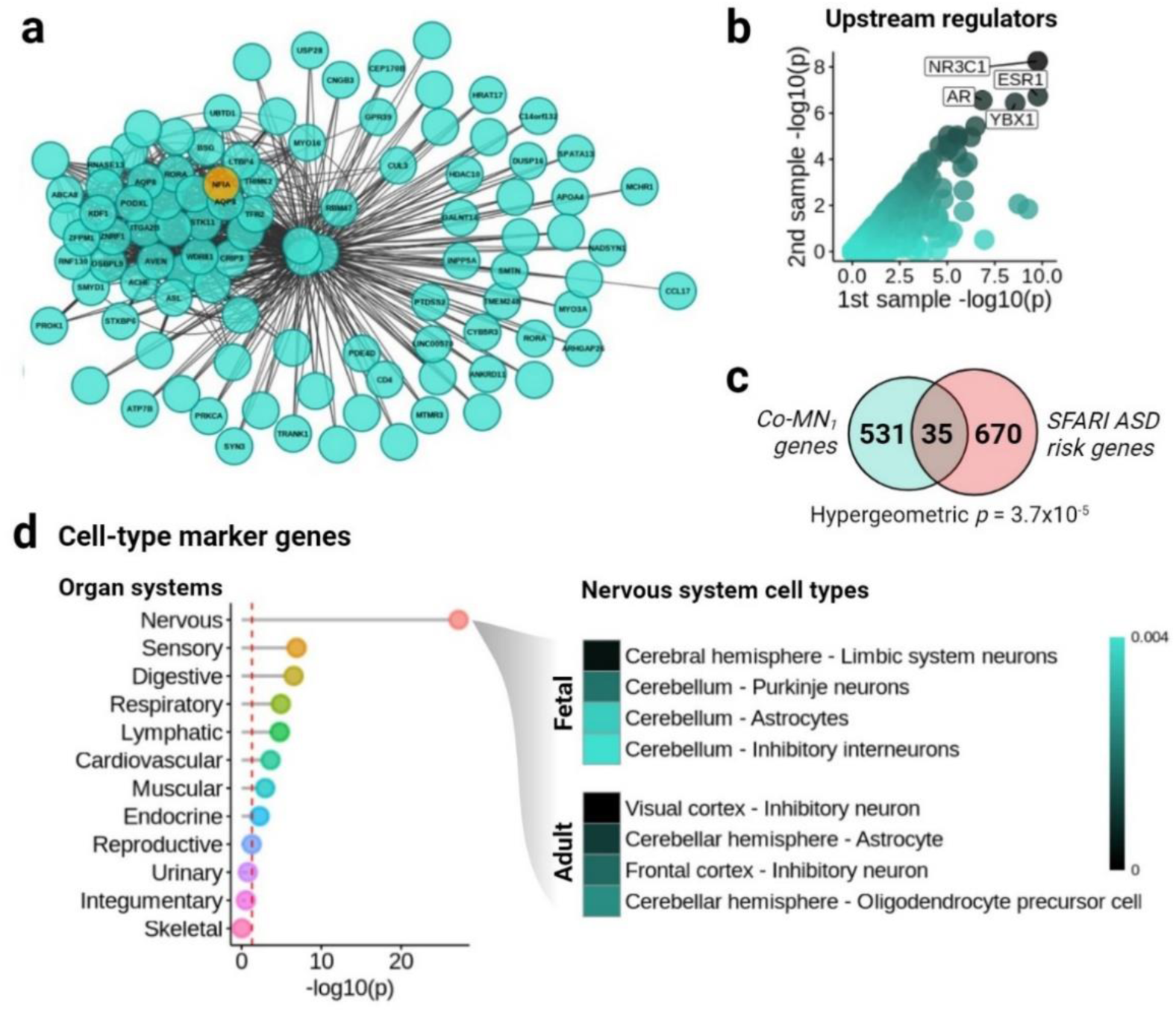
Biological interpretation of key co-methylation network, Co-MN_1_, mediating the effect of prenatal DEHP exposure on ASD and ADHD symptoms. **a** *Visualisation of Co-MN_1_*. Nodes are CpG sites and edges represent correlations. Shown are the top 100 most intercorrelated CpG sites and their adjacent genes. The most highly connected CpG site in Co-MN_1_ (mapping to *NFIA*) is highlighted in orange. b, c, d *Enrichment analyses of Co-MN_1_’s 531 genes*. **b**: Upstream regulators of Co-MN_1_’s gene set inferred using the LISA platform. LISA provides p-values for each regulator across five sample types (see Methods); the two lowest p-values are shown for each regulator. Labelled are the top four of 1316 regulators tested in total. **c**: *Enrichment of Co-MN_1_ for ASD risk genes from the SFARI database.* d: Enrichment of Co-MN_1_ for fetal and adult cell-type marker genes, with p-values (left) aggregated at the organ-system level using Fisher’s method, and (right) shown at the cell-type-specific level within fetal and adult nervous-system tissue. Markers of 1355 distinct cell types were tested in total.

#### 2.6.1 Hormone-receptor targets

We tested for upstream regulators likely controlling Co-MN_1_ by analysing the 531 genes using LISA.^37^ In brief, LISA identifies upstream regulators based on the enrichment of their targets in the input gene set.^37^ Since DEHP is an endocrine disruptor, evidence of upstream hormone signalling was of particular interest.^38^ Strikingly, among 1316 regulators tested, the two most significant were indeed endocrine-related (Figure 4b): estrogen receptor-alpha (ERα; encoded by the gene *ESR1*) (*p* = 1.7×10^-10^) and the glucocorticoid receptor (GR; encoded by *NR3C1*) (*p* = 1.8×10^-10^). Ranked fourth was the androgen receptor (AR, encoded by *AR*) (*p* = 1.4×10^-7^). DEHP is known to interfere with the function of all three receptors.^39,40^

#### 2.6.2 Neural cell-type markers

Given its association with neurodevelopmental outcomes, it was plausible that Co-MN_1_ could represent, in cord blood, an epigenetic memory of alterations more relevant to other, especially neural, tissue types. To elucidate the tissue types and developmental stages likely most impacted by disruption of Co-MN_1_, we analysed the gene set for signatures of 1355 foetal and adult cell types using WebCSEA (Figure 4d).^41^ The network was most strongly enriched for nervous-system cell-type marker genes (adjusted Fisher’s method *p* = 6.2×10^-28^), followed by sensory (*p* = 1.26×10^-7^) and digestive (*p* = 2.92×10^-7^) systems. We next investigated which individual fetal and adult nervous-system cell types were most enriched. At the fetal stage, the highest ranked cell types were cerebral limbic-system neurons (*p* = 0.0007), cerebellar Purkinje neurons (*p* = 0.002) and cerebellar astrocytes (*p* = 0.003). In adult tissue, the highest ranked cell types were visual cortex inhibitory neurons (*p* = 0.0004), cerebellar astrocytes (*p* = 0.001) and frontal cortex inhibitory neurons (*p* = 0.002). In contrast, no enrichment was observed for any cord blood cell-type signatures (p-value range: 0.36 to 0.97). These findings suggest that Co-MN_1_ more closely encapsulates neural than cord-blood-related epigenetic processes.

#### 2.6.3 ASD risk genes

If, as our findings indicated, Co-MN_1_ mediates the effect of the prenatal environment on ASD and ADHD symptoms, it might be expected to show enrichment for known risk genes for these outcomes. Examining ASD risk genes from the Simons Foundation Autism Research Initiative (SFARI) database^42^ (Figure 4c), we found a significant overlap that included the transcription factors *FOXP1* and *RORA* (a key regulator of brain estrogen levels,^43^) the synaptic scaffolding gene *SHANK2*, and the axon-guidance genes *SEMA5A* and *PLXNB1* (overall, 35 of 670 SFARI genes here present, significantly more than expected by chance, hypergeometric *p* = 3.7×10^−5^).

A previous study by Nardone *et al.*^44^ used WGCNA to identify two Co-MNs associated with ASD diagnosis in postmortem brain tissue (labelled “green-yellow” and “brown” in the study, respectively). To explore whether these ASD-associated networks were also related to Co-MN_1_, we tested the intersection between the 531 genes in Co-MN_1_ and the genes in these networks. A modest but highly significant overlap with both networks was evident (23 of 309 “green-yellow” genes captured, hypergeometric p = 3.6×10^-6^; 35 of 577 “brown” genes captured, hypergeometric *p* = 1.5×10^-6^; 57 genes shared with both networks in total, hypergeometric *p* = 6.8×10^-11^).

In summary, these additional bioinformatics analyses highlight the biological plausibility of Co-MN_1_ as a mediator linking DEHP to ASD and ADHD symptoms, suggesting a role in the development of key components of the nervous system, including the cerebellum, under the control particularly of estrogen, glucocorticoid and androgen signalling.

### 2.7 Co-methylation network Co-MN_1_ is associated with elevated inflammation at birth

Neurodevelopmental conditions are often associated with elevated inflammation.^45^ In this cohort, higher inflammation at 28 weeks gesation and at birth is associated with adverse neurodevelopmental outcomes.^46–48^ To further validate Co-MN_1_ as a signature of disrupted neurodevelopment, we investigated its relationship with the inflammatory marker glycoprotein acetyls (GlycA) in cord blood.

Co-MN_1_ showed a strong and positive correlation with birth GlycA levels (*ρ*= 0.19, p = 2.6×10^-8^) (Figure 3e), a finding confirmed by adjusted linear regression (*β* = 0.37, p = 0.033). Although prenatal DEHP exposure is not associated overall with birth GlycA levels,^49^ we hypothesised that DEHP’s perturbation of Co-MN_1_, specifically, might elevate GlycA levels. Mediation analysis confirmed this hypothesis (*β_ACME_* = 0.011 [95% CI 0.004, 0.019], *p* = 0.004) (Figure 3f).

### 2.8 Co-methylation network Co-MN_1_ also mediates the neuroprotective effects of a maternal wholefoods diet

We next reasoned that if Co-MN_1_ is a systems-level mechanism underlying ASD and ADHD symptoms, mediating the effects of DEHP exposure on these neurological phenotypes, it might also mediate the effects of other exposures. To investigate the broader relevance of Co-MN_1_ beyond DEHP, we selected maternal prenatal wholefoods diet as an additional exposure. This dietary pattern is associated with improved neurodevelopmental outcomes in BIS,^46^ consistent with other studies that have shown a relationship between a maternal wholefoods diet and reduced ASD and ADHD symptoms.^50^ We investigated whether a maternal wholefoods diet reduces ASD and ADHD symptoms in part by altering methylation across Co-MN_1_ (Figure 3d). Substituting the wholefoods diet measure^46^ (detailed in Methods) for DEHP as exposure in the mediation models, we observed a striking reversal of effect directions, with a higher wholefood diet associated with increased SDQ_4yrs_ prosocial behaviour (*β_ACME_* = 0.019, p = 0.036, proportion mediated = 0.62), and reduced CBCL_2yrs_ attention problems (*β_ACME_* = - 0.022, p = 0.048, proportion mediated = 0.20) (Figure 3f). Through Co-MN_1_, maternal wholefoods diet also showed an opposite effect on birth GlycA levels, with a significant reduction evident (*β_ACME_* = −0.003, p < 0.0001, proportion mediated undefined)(Figure 3f).

Intriguingly, prenatal DEHP exposure and maternal wholefoods diet are positively correlated (*ρ* = 0.12, *p* = 0.001). Their opposing effects on ASD and ADHD symptoms (via Co-MN_1_) therefore cannot be explained by the hypothesis that DEHP and wholefoods diet are merely (inversely related) proxies for each other or a common antecedent factor.

### 2.9 The two DEHP-associated epigenetic signatures, MPS_DEHP_ and Co-MN_1_, are associated with ASD across independent blood and postmortem-brain DNA methylation datasets

*2.9.1 Independent blood and postmortem brain DNA methylation datasets*

To validate our cord blood–derived findings and confirm their relevance to DNA methylation in brain tissue, we investigated whether the associations between both MPS_DEHP_ and Co-MN_1_ and adverse neurodevelopmental outcomes would replicate in two external Illumina 450k DNA methylation datasets (Figure 5a):

1. **Whole blood** (n = 66 ASD, ages 3–9 years, mean of 6 years) with a severity scale measured using the Childhood Autism Rating Scale (CARS).^51^
2. **Postmortem brain** (n = 40; 21 ASD, 19 controls; ages 2–56 years, mean of 25.1 years) with methylation profiled across the prefrontal and temporal cortices and the cerebellum.^16^

**Fig. 5.**
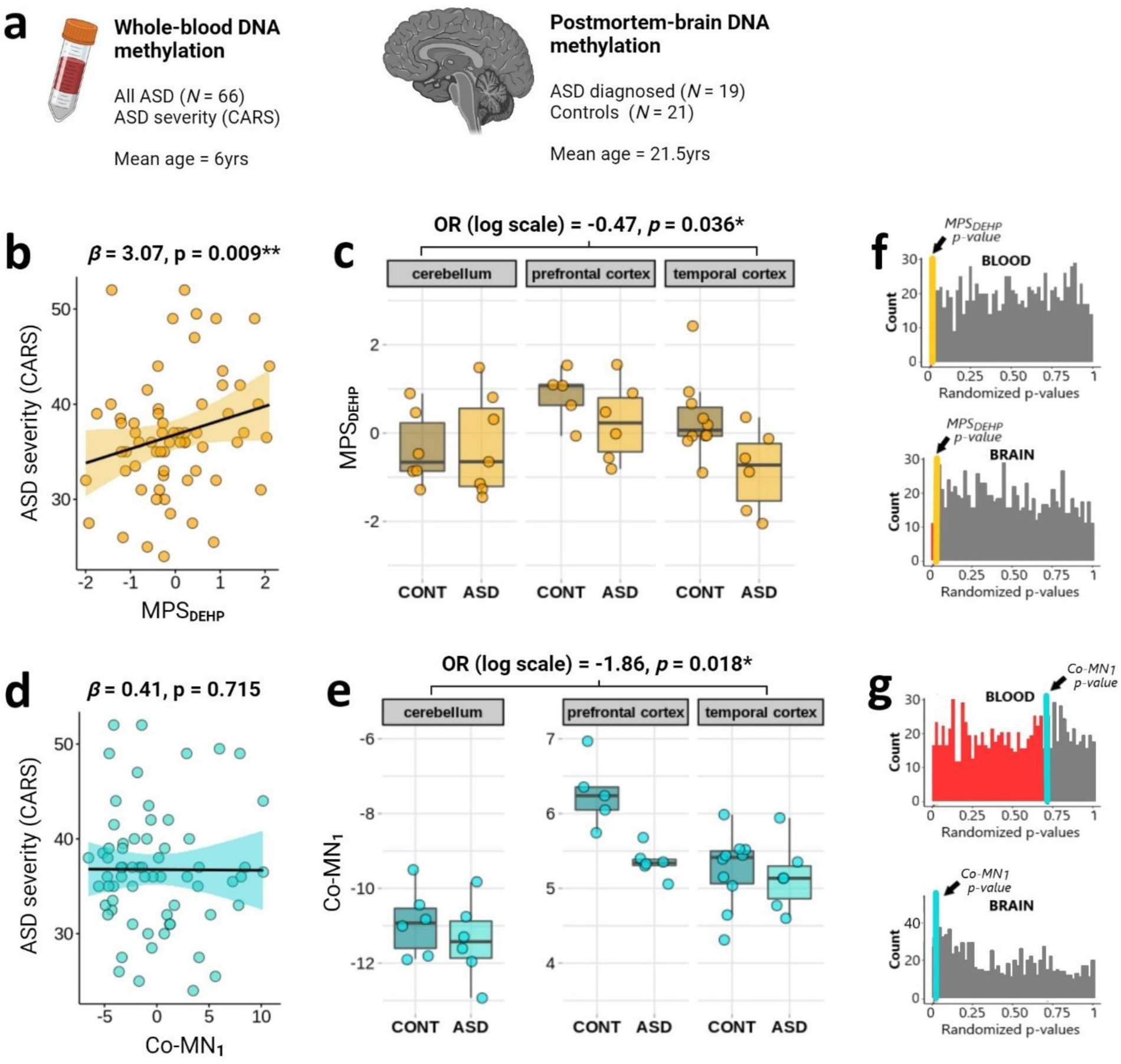
Independent replication of association between DEHP-related epigenetic factors and ASD. **a** We reconstructed MPS_DEHP_ and Co-MN_1_ in independent blood (left) and postmortem brain (right) DNA methylation datasets and assessed their association with ASD severity and diagnosis, respectively. **b** Association between MPS_DEHP_ and ASD severity in peripheral blood. **c** Association between MPS_DEHP_ and ASD diagnosis (case vs. control (CONT)) in postmortem brain (across the three regions combined). **d** Association between Co-MN_1_ and ASD severity in peripheral blood. **e** Association between Co-MN_1_ and ASD diagnosis in postmortem brain (across the three regions combined). **f, g** Distribution of p-values for the association between randomised MPS and Co-MNs and ASD severity (in blood) or diagnosis (in brain) (the order corresponds to plots a-d). The coloured lines indicate, for comparison, the actual p-values for MPS_DEHP_ (gold) and Co-MN_1_ (turquoise). Almost all randomised MPS and Co-MNs show no association with ASD, demonstrating that the significant replication is not due to hidden bias.

*2.9.2 Replication analysis of methylation profile score*

We first reconstructed MPS_DEHP_ in both datasets by summing methylation for each participant across the DEHP-associated CpG sites, weighted by their effect estimates from BIS. Given the sparser coverage of the 450k array compared to the EPIC 850k array used in BIS, for replication analyses, we used only the highest-confidence 150 CpG sites available (ranked by significance for DEHP in BIS) to avoid possible bias towards lower-confidence CpG sites.

We then examined the association between MPS_DEHP_ and ASD. In whole blood, MPS_DEHP_ was strongly and positively associated with ASD severity (*β* = 3.07, p = 0.009), consistent with its relationship in cord blood to ASD and ADHD symptoms in BIS (Figure 5b). In postmortem brain, MPS_DEHP_ was significantly associated with ASD diagnosis but with a negative effect direction (OR [log scale] = −0.47, p = 0.036; Figure 5c).

#### 2.9.3 Replication analysis of co-methylation network

Similarly, we reconstructed Co-MN_1_ in the whole blood and postmortem brain datasets using the highest-confidence 150 CpGs from the network available in each dataset (ranked by intercorrelation within Co-MN_1_ in BIS). In whole blood, Co-MN_1_ (represented using its first principal component) was not associated with ASD severity (*β* = 0.41, p = 0.715; Figure 5d). In postmortem brain, Co-MN_1_ was strongly and negatively associated with ASD diagnosis (OR [log scale] = −1.86, p = 0.018; Figure 5e), consistent with MPS_DEHP_’s effect direction.

The presence of shared epigenetic signatures in blood and brain with opposite effect directions was not unexpected, aligning with cross-tissue analyses of neurological disorders. For example, a recent meta-analysis of DNA methylation changes in Alzheimer’s disease found 97 CpG sites significantly associated with case status in both blood and brain, and for 74 of these CpG sites the effect directions were opposite.^52^

#### 2.9.4 Calculation of empirical p-values to confirm replication

To ensure that these results were not influenced by systematic bias, we evaluated how frequently MPSs and Co-MNs constructed from random sets of 150 CpGs were associated with ASD in the replication datasets. If systematic bias were present, a significant proportion of randomised MPS and Co-MNs should show a comparable association with ASD. However, of 1000 randomised MPS, p-values for only 0.6% in whole blood (against ASD severity) and 3% in postmortem brain (against ASD diagnosis) were lower than those for MPS_DEHP_. Using these frequencies, we calculated empirical p-values for MPS_DEHP_, which were *p* = 0.006 in blood and *p* = 0.030 in brain (Figure 5f), closely matching the regression p-values for MPS_DEHP_ in these datasets. Similarly, comparing Co-MN_1_ against 1000 randomised Co-MNs, we found that 69.7% of p-values for the randomised Co-MNs in blood and 2.7% in postmortem brain were lower than those for Co-MN_1_, giving empirical p-values of *p* = 0.696 and *p* = 0.027, respectively (Figure 5g), again matching the corresponding regression results. These findings indicate that the observed replication is not due to study-specific bias and that MPS_DEHP_ and Co-MN_1_ are genuine epigenetic signatures of adverse neurodevelopment.

## 3 Discussion

Here we employed systems-biology approaches in a large population-based cohort to investigate the role of cord blood-measured DNA methylation in mediating the association between prenatal DEHP exposure and ASD and ADHD symptoms at ages 2 and 4 years. Compelling evidence was observed for mediation through both (1) a methylation profile score for DEHP exposure (MPS_DEHP_) and (2) a brain-related network of co-methylated CpG sites (Co-MN_1_). Both the MPS_DEHP_ to ASD and the Co-MN_1_ to ASD associations were replicated in independent blood and postmortem brain DNA methylation datasets, validating MPS_DEHP_ and Co-MN_1_ as signatures of adverse neurodevelopment.

High-dimensional molecular epidemiological datasets allow the application of statistical and computational techniques that make causal inference possible even in an observational setting.^26,53^ Here, the use of multiple complementary approaches allowed us to triangulate evidence on the key finding that DEHP exposure is associated causally with ASD and ADHD symptoms via epigenetic changes.

Using MPS_DEHP_, we demonstrated that broad changes in cord-blood DNA methylation elicited by DEHP exposure were not only associated strongly with ASD and ADHD symptoms, but mediated the effects of prenatal DEHP exposure on these symptoms. Methylation profile scores have been used to summarise DNA methylation changes associated with various conditions, such as increasing age^54^ and exposures such as smoking and alcohol consumption.^55,56^ By aggregating effects across many differentially methylated CpG sites (in this case 1000) into a single score per participant, MPSs can improve statistical power to detect and replicate subtle effects that are distributed throughout the epigenome. While DEHP’s effects on individual CpG sites may be small or present only in subgroups of samples, MPS_DEHP_ summarised these CpG-specific effects across the cohort. This provided a clearer epigenetic signature of prenatal DEHP exposure to analyse for mediated effects. Aggregate approaches can also increase discovery power by limiting the multiple testing burden.

The identification of a co-methylation network, Co-MN_1_, as an alternative systems-level mediator of DEHP’s effects on ASD and ADHD symptoms sheds light on the likely mechanisms involved. Consistent with DEHP’s endocrine-disrupting properties, Co-MN_1_ was enriched for targets of hormone receptors with which DEHP is known to interfere. These included the estrogen receptor (ERα, encoded by the gene *ESR1*, ranked 1^st^), the glucocorticoid receptor (GR, encoded by *NR3C1*, ranked 2^nd^) and the androgen receptor (AR, encoded by *AR*, ranked 4^th^),^39,40^ all of which can recruit epigenetic agents, including DNA methyltransferases (DNMTs), to alter methylation of target genes.^57^ Of particular interest, the most interconnected CpG within Co-MN_1_ was located within the *NFIA* gene, which is itself regulated by estrogen and glucocorticoid signalling^58^ and is critical in neurodevelopment.^58,59^ For instance, mutations in *NFIA* can cause NFIA-related disorder, characterised by corpus-callosum malformation and ASD and ADHD symptoms.^35^

Further underscoring its biological plausibility, Co-MN_1_ was enriched for ASD risk genes from the SFARI database and showed a substantially stronger overrepresentation of nervous-system cell-type marker genes than those of any other organ system. Consistent with this, despite the cord blood origin of the DNA methylation data, Co-MN_1_ showed no enrichment for cord blood cell-type markers, suggesting that Co-MN_1_ is not a cord blood-related mechanism *per se*. Moreover, most SFARI ASD risk genes are derived from genetic studies. The fact that, here, DEHP exposure is associated with epigenetic disruption of genes that themselves show genetic—and therefore arguably causal—links to ASD provides additional evidence that the association between DEHP exposure and ASD, through Co-MN_1_, is causal in nature.

The specific fetal and adult neural cell types identified are of interest. Most enriched at the fetal stage were markers of cerebral (limbic system) and cerebellar (Purkinje neuron, astrocyte, and inhibitory interneuron) cell types. The limbic system has well-established links to ADHD, playing key roles in emotional and behavioural responses and attention in conjunction with higher-order control centres in the prefrontal cortex.^60^ Likewise, while the cerebellum historically was associated primarily with the calibration of motor output, it is now known to play related roles in fine-tuning social and emotional behaviour.^61^ For example, the cerebellum is integral to “theory of mind”, or the ability to attribute mental states to oneself and others, a crucial aspect of social cognition.^61,62^ Indeed, cerebellar abnormalities have been identified in MRI analyses of both prenatal DEHP exposure and, separately, ASD diagnosis, as well as in rodent studies of DEHP exposure.^63,64^ It is also noteworthy that while hormonal signalling is essential across the developing brain, *ESR1* and *NR3C1* show especially high expression in limbic system structures and the cerebellum (Supplementary Figure 4 from the Human Protein Atlas^65^), perhaps entailing increased vulnerability in these regions to DEHP exposure.

At the adult developmental stage, Co-MN_1_’s enrichment for cerebellar cell-type marker genes remained strong, with cerebellar astrocytes and cerebellar oligodendrocyte precursor cells among the top four findings. Markers of adult visual cortex inhibitory neurons were also strongly enriched. Although the visual cortex is distal to primary brain regions associated with executive control and social behavior, alterations in its structure and differences in visual information processing are well-documented in ASD and ADHD.^66–69^ Additionally, adult frontal-cortex inhibitory neuron markers were highly enriched, aligning with the fetal-tissue findings for the limbic system, which, as noted, is intricately connected to frontal executive-control centers.^60^ This pattern is further supported by the replication results in the postmortem-brain dataset, which primarily included adult samples and in which Co-MN_1_ was most strongly associated with ASD diagnosis in the frontal cortex (Figure 5e).

Of note, our finding that a maternal wholefoods diet exerts an apparent neuroprotective effect through Co-MN_1_ suggests that this network may be a convergent axis for multiple factors involved in the aetiology of ASD and ADHD rather than a DEHP-specific module. Maternal diet has well-established links to offspring neurodevelopment. For instance, a recent study involving two large prospective European cohorts, the Norwegian Mother and Child Cohort Study (MoBa) and the Avon Longitudinal Study of Parents and Children (ALSPAC), found that a healthy maternal diet was associated with a reduced likelihood of child ASD diagnosis and reduced social communication difficulties.^50^ Co-MN_1_ provides an epigenetic mechanism that may partly explain these associations while also highlighting the potential for dietary interventions during pregnancy to mitigate the adverse effects of DEHP exposure.

Replication of the MPS_DEHP_ and Co-MN_1_ to ASD associations in independent blood and postmortem brain DNA methylation datasets provided further evidence that these factors— disrupted epigenetically by DEHP exposure—are *causally* linked to neurodevelopmental outcomes. Postmortem-brain replication also points to the utility of cord blood, accessible non-invasively in human cohorts, in elucidating DNA methylation changes in the developing brain through possible epigenetic memories preserved across tissue types. Nonetheless, the observed differences in effect direction between blood and brain highlight the need for caution when interpreting directionality if blood is used as a proxy for brain. These differences align with findings from cross-tissue analyses of other brain-related disorders and may reflect tissue-specific activation or inhibition of hormone pathways in response to DEHP exposure.^52^ Even between neurons and glial cells in rodents, DEHP has been found to produce opposing effects on expression of Cyp1A1 and other genes.^70^

Importantly, in this cohort, where causal mediation was demonstrated using the continuous DEHP exposure measure, mean DEHP exposure (1.6 μg/kg bw/day) is well below the current tolerable daily intake (TDI) in the European Union (50 μg/kg bw/day),^71^ suggesting that estimated safe levels require reassessment. This is already occurring in some countries. For example, Norway has recently included three specific phthalate chemicals in its list of Substances of Very High Concern for Authorisation, to be phased out entirely.^72^ Tighter phthalate regulations are also planned globally in response to mounting evidence of harm.^72^ Existing standards are particularly concerning given that safe levels are generally estimated for DEHP in isolation, while real-world exposures involve chemical mixtures with a cumulative effect likely to be substantially stronger than that of DEHP alone.^73^

A key strength of this study is the use of systems-biology approaches to identify broader convergent epigenetic alterations linking prenatal DEHP exposure toASD and ADHD symptoms. The availability of DEHP-exposure measures, DNA methylation profiling, and neurodevelopmental assessments on the same participants in the BIS cohort allowed us to employ formal mediation analysis to test for causal effects. Extensive measures on each participant allowed systematic screening and adjustment for potential confounding variables in the mediation models. Several lines of evidence supported the central findings from this causal inference technique. First, **temporality**: the prospective mediation analyses confirmed the sequence of DEHP exposure (prenatal), DNA methylation changes (at birth), and neurodevelopmental outcomes (postnatal), fulfilling the essential requirement that cause precedes effect. Second, **consistency**: DEHP showed mediated effects through (a) two epigenetic factors, MPS_DEHP_ and Co-MN_1_, identified using independent methods, to (b) multiple ASD and ADHD symptoms scales, and at (c) two time points, reducing the likelihood of spurious effects. Third, **coherence** and **biological plausibility**, which Co-MN_1_ showed across multiple domains, including enrichment for neural cell type markers and for genes linked genetically and therefore causally to ASD. An established neuroprotective exposure, prenatal maternal wholefoods diet, also operated through Co-MN_1_, in an opposing direction. Lastly, **replicability**: MPS_DEHP_ and Co-MN_1_ were associated with adverse neurodevelopment across three distinct and independent datasets, including ASD diagnosis (not merely symptoms) in postmortem brain tissue.

Several limitations should be noted. Causal mediation analysis could only be applied in BIS, as direct DEHP-exposure measures were unavailable in the replication datasets. While we present causal evidence, this study is not interventional, and ethical considerations prevent such a design in human studies. However, in future studies we plan to revisit key findings, including to further develop and validate MPS_DEHP_, using controlled animal experiments and/or 3D human brain organoids.

In summary, this study suggests that prenatal DEHP exposure increases symptoms of ASD and ADHD through systems-level changes in DNA methylation. Computational analyses indicate that these epigenetic alterations are driven by disruptions in hormone-signalling pathways and differentially impact the development of the cerebellum, limbic system and frontal cortex—brain regions crucial for social behaviour and attentional control. These findings provide causal insights into DEHP’s effects on the developing brain and reinforce growing concerns regarding the risk of exposure during key early neurodevelopmental windows.

## 4 Methods

### 4.1 Study cohort

The BIS is a population-derived birth cohort from Victoria, Australia, comprising 1074 mother-infant pairs and aimed at investigating the early-life causes of non-communicable diseases.^74^ Women were recruited from 2010 to 2013 between 15 and 28 weeks of completed pregnancy but subsequently excluded if their child was born before 32 weeks or diagnosed with a congenital disorder or a serious illness. As described elsewhere, extensive biological, clinical, and questionnaire measures were collected prenatally, at birth, and in intervals up to age 9 years.^74^ The study was approved by the Barwon Health Human Research Ethics Committee (HREC 10/24), with written informed consent obtained from the participating families.

### 4.2 Prenatal DEHP exposure

Phthalate metabolite levels were measured in 842 pregnant women using a single-spot urine specimen collected at 36 weeks of gestation. High-performance liquid chromatography– tandem mass spectroscopy with direct injection was conducted by the Queensland Alliance for Environmental Health Science (QAEHS). Their procedures are detailed elsewhere.^75^ For monoethyl phthalate (MEP), monoisobutyl phthalate (MiBP), mono-n-butyl phthalate (MnBP), mono-(2-ethyl-5-hydroxyhexyl) phthalate (MEHHP), and mono-(2-ethyl-5-oxohexyl) phthalate (MEOHP), repeated spot specimens collected in the third trimester had intra-class correlation coefficients over 0.4 in one or both of two previous studies.^76,77^ This indicates that single-spot tests can capture third-trimester phthalate exposure with reasonable reliability.

Phthalate metabolite measurements were corrected for batch, specific gravity and time of day of sample collection.^10^ Phthalate estimated daily intake was then computed accounting for maternal prenatal weight, fractional excretion of the compound, and compound-to-metabolite molecular weight ratio.^10^ The metabolites MEHHP, MEOHP and mono-(2-ethyl-5-carboxypentyl) phthalate (MECPP) were used to calculate di-(2-ethylhexyl) phthalate (DEHP) daily intake. As in our previous work,^10,30^ phthalate measures were logarithm base 2 transformed for analyses.

A high DEHP-exposure variable, DEHP_HIGH_, was defined using the top 2% of participants with DEHP measured vs. the rest.

### 4.3 ASD and ADHD outcomes

ASD and ADHD symptoms were assessed at 2 years of age using the Child Behavior Checklist for ages 1.5–5 (CBCL_2yrs_)^28^ and at 4 years of age using the Strengths and Difficulties Questionnaire P4-10 (SDQ_4yrs_).^29^ Both are widely-used parent-reported questionnaires. The CBCL consists of 99 items scored on a three-point Likert scale (0, not true; 1, somewhat or sometimes true; or 2, very true or often true). The CBCL has shown high test-retest consistency,^28,78^ and parent-reported scores correlate strongly with direct measures of child neurobehaviour.^79^ Here the DSM-5-oriented *autism spectrum problems, attention problems*, and *attention-deficit/hyperactivity problems* subscales of the CBCL_2yrs_ were used. The SDQ consists of 25 items, also scored on a three-point Likert scale (0, not true; 1, somewhat true; or 2, certainly true) with validated psychometric properties.^80^ The SDQ_4yrs_*peer problems*, *prosocial behavior,* and *hyperactivity* subscales were used.

In this cohort, CBCL_2yrs_ *autism spectrum problems* predicted subsequent doctor-diagnosed autism with an area under the curve (AUC) of 0.92.^48^ In the literature, SDQ_4yrs_ peer problems has moderate to high accuracy in distinguishing preschoolers with ASD from those who are developmentally typical (AUC 0.82).^81^ Moreover, symptom scores have the advantage that they reflect the continuous nature of ASD and ADHD phenotypes and are sensitive to phenotypes present before the typical age of diagnosis (mean ASD diagnosis age is 5.9 years in Australia).^82^

### 4.4 Whole genome DNA methylation profiling and genotyping

The Illumina Infinium MethylationEPIC BeadChips (referred to henceforth as “EPIC array”) was used for DNA methylation profiling of cord blood from the BIS cohort. Genomic DNA (200 to 500 ng) from cord blood was randomised into 96-well plates and sent to the Kobor laboratory (University of British Columbia, Canada) for sodium bisulfite treatment and processing on the EPIC array. The EPIC array measures the DNA methylation level at more than 850, 000 CpG sites (referred to as “EPIC probes”), and covers all gene promoters, gene bodies and ENCODE-assigned distal regulatory elements.^83^ Raw IDAT files were processed and analysed using the MissMethyl and minfi packages for R.^84,85^ Samples were checked for quality and those with a mean detection p-value of >0.01 were removed (128 samples), leaving 946 cord blood samples for analysis. Data were normalised for both within and between array technical variation using SWAN (Subset-quantile Within Array Normalization).^86^ Probes with poor average quality scores (detection p-value > 0.01) and cross-reactive probes^84^ were removed from further analysis. This left a total of 798,259 probes for analysis. Cell composition was determined using the *estimateCellCounts* tool, with the “CordBlood” reference data used for neonatal blood spot analysis.^87^ Maternal contamination of infant cord blood was determined using a CpG signature, with maternally contaminated samples being removed from the epigenetic analyses.^88^ Whole-genome genotyping was also performed using DNA samples extracted from cord and 12-month whole blood (see Supplementary Methods).

### 4.5 Other measures

Maternal wholefoods dietary patterns were derived using principal component analysis as described previously.^89^ Principal component 1 (PC1) captured a wholefoods dietary pattern, with high loadings of fish, nuts, eggs, green vegetables, and whole grains.

### 4.6 External replication datasets

The datasets used for replicating the associations between DNA methylation and neurodevelopmental outcomes are described in detail elsewhere.^51,16^ Briefly, DNA methylation in both datasets was profiled using the Illumina 450k array. After pre-processing, the whole-blood dataset consisted of 428, 560 CpG sites from 66 ASD-diagnosed participants from Brazil (ages 3-9 years (mean = 6 years)), each scored for ASD severity using the Childhood Autism Rating Scale (CARS).

The postmortem brain dataset consisted of 428, 526 CpG sites profiled on 19 ASD cases and 21 controls from the USA (ages 2-56 years (mean = 25.1 years)), with samples taken in separate participants from one of three brain regions: frontal cortex (n = 11), temporal cortex (n = 16), and cerebellum (n = 13).

### 4.7 Statistical analysis

#### 4.7.1 Epigenome-wide association study for DEHP exposure

To screen for DEHP-associated CpG sites, an EWAS was conducted for DEHP_HIGH_ using the missMethyl R package.^84^ The models were fit using limma, with methylation M values as the dependent variable, adjustment for age and sex, and `robust=TRUE` to handle CpG outliers. Within the missMethyl pipeline, the RUV-III algorithm was applied to remove variation from unwanted sources, such as unmeasured processing factors, using a threshold of p > 0.9. Full covariate adjustment was performed during formal mediation analysis to allow nuanced handling of possible confounders. CpG sites were ranked by statistical significance for DEHP_HIGH_ and high-ranking CpG sites were analysed for potential mediators, as outlined below, using a methylation profile score and a co-methylation network analysis.

#### 4.7.2 Methylation profile score for DEHP exposure

A methylation profile score for prenatal DEHP exposure was calculated by summing the methylation M values, *m*, for the top *n* = 1000 DEHP_HIGH_-associated CpGs weighted by their EWAS effect estimates, *w*:

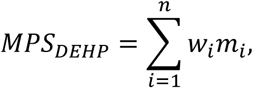

#### 4.7.3 Co-methylation network analysis

Co-MNs were identified within the top 2000 DEHP_HIGH_-associated CpGs using WGCNA,^34^ An unsigned weighted adjacency matrix was first constructed from the 2000 CpGs using a soft-thresholding power of 4 to approximate a scale-free network (R^2^ = 0.8). The adjacency matrix was then transformed into a Topological Overlap Matrix (TOM) to account for indirect connections between CpGs (i.e. via “neighbour” CpGs). In the resulting network, nodes represent CpGs and edges reflect co-methylation relationships between CpGs.

Co-MNs were next identified using hierarchical clustering followed by a dynamic tree-cut algorithm, with a minimum cluster size of 15 and merge height of 0.25. For statistical analysis, each Co-MN was summarised using its first principal component.

Spearman correlation was used to screen for potential mediators. Co-MNs correlating (p < 0.1) with DEHP exposure and at least one ASD- or ADHD-related outcome were taken to causal mediation analysis.

#### 4.7.4 Causal mediation analysis

Causal mediation analysis was conducted using the `mediation` R package.^33^ This procedure partitions the total effect of an exposure E on an outcome O, estimated here using adjusted linear regression, into its (i) *indirect effect*, operating causally via the putative mediator M, and (ii) *direct effect*, operating via other mechanisms. Bootstrapping was employed to calculate stable confidence intervals. Estimated proportions mediated were not reported if they fell outside the range of a true proportion, [0,1]. This can occur if the direct effect is small (and the indirect or mediated effect is large) or direct and indirect effect estimates have opposite signs. Indirect effect p-values were not adjusted for multiple comparisons across the six neurodevelopmental outcomes because these outcomes are not independent, as assumed by the Bonferroni and Benjamini-Hochberg procedures, but rather capture different components of the ASD and ADHD phenotypes (which are themselves related). An important consequence of this dependence, however, is that a true signal should manifest in consistent effects across multiple outcomes, reflecting their shared biological underpinnings.

#### 4.7.5 Covariate adjustment

A key assumption of this causal-mediation framework is the absence of unmeasured confounding in the E-O (total effect), E-M and M-O relationships. A literature-based and data-adaptive epidemiological approach was applied to refine a covariate set for each model.^53^ All models were adjusted for age and sex, as well as principal component 1 of the BIS genotyping dataset to account for genetic architecture. E-O and M-O models were adjusted for covariates relevant to the CBCL and SDQ neurodevelopmental outcomes, including age at review. Since the initial DEHP_HIGH_ EWAS was minimally adjusted, E-M models were adjusted for cell composition using the following seven cord blood cell-type proportions unless a given cell-type proportion showed evidence of itself mediating the E-M relationship and so lying on the causal pathway:^90^ CD8+ T cells, CD4+ T cells, natural killer (NK) cells, B cells, monocytes, granulocytes, and nucleated red blood cells (nRBCs). For the causal mediation analyses of prenatal DEHP exposure, all seven cell type proportions were included in the models. For the causal mediation analyses of maternal wholefoods diet, granulocytes and nRBCs were excluded as they partly mediated the effect of diet on DNA methylation. As in previous work,^46^ additional potential confounders were determined through a data-driven screening procedure across 60 relevant pre- and perinatal environmental factors (see Results for full lists of covariates included).

#### 4.7.6 Functional enrichment analysis

To facilitate biological interpretation, CpG sites within any Co-MN demonstrating causal mediation were annotated to their corresponding genes using UCSC (missMethyl default) or Ensemble^91^ (for key CpGs missing from UCSC annotation), with Genome Build 37 (GRCh37). The resulting gene set was investigated using bioinformatics analyses. The *Epigenetic Landscape In Silico Deletion Analysis (LISA*) platform was used to infer potential transcription factors and upstream regulators.^37^ LISA infers upstream regulators across five different sample types; inferred regulators were filtered to retain only those that achieved statistical significance in at least two out of the five analyses. *Web-based Cell-type-Specific Enrichment Analysis of Genes (WebCSEA)* was used to characterise affected cell, tissue and organ types at both fetal and adult developmental stages.^41^ Fisher’s method was employed to calculate a p-value per organ system from the cell-type-specific p-values provided by *WebCSEA*, and the resulting p-values—adjusted for multiple comparisons using the Benjamini-Hochberg procedure—were used to provide a ranking of organ systems. Both *LISA* and *WebCSEA* account for potential gene-length bias. The hypergeometric test was applied to test for overlap with (i) ASD risk genes from the SFARI database,^92^ and (ii) a gene set derived from a previously published *WGCNA* co-methylation network analysis of ASD diagnosis using postmortem human brain tissue.^44^

#### 4.7.7 Replication analyses

MPS_DEHP_ was reconstructed in the whole-blood and postmortem brain DNA methylation datasets by summing methylation M values across the top 150 CpG sites from the DEHP_HIGH_ EWAS in BIS weighted by their effect estimates from this EWAS. Co-MN_1_ was reconstructed in each dataset using the top 150 available CpGs ranked by intercorrelation within Co-MN_1_ in BIS. The association between each epigenetic variable and ASD severity in the whole-blood dataset was assessed using linear regression. Models were adjusted for age, sex, and principal component 1 of the methylation dataset. In postmortem brain, the association between each epigenetic variable and ASD diagnosis was assessed across the prefrontal cortex, temporal cortex, and cerebellum jointly using logistic regression adjusted for age, sex, brain region and methylation principal component 1.

A corresponding empirical p-value was calculated for each association by (i) generating 1000 random sets of 150 CpG sites (sampling from the 450k array), (ii) constructing from each set a randomised MPS (when assessing MPS_DEHP_ models) or taking principal component 1 (when assessing Co-MN_1_ models), (iii) testing the association between the randomised epigenetic factor and the outcome of interest (ASD severity or diagnosis), and (iv) calculating the proportion of randomised factors with a lower p-value *p*_rand_ than MPS_DEHP_ or Co-MN_1_. The empirical p-value for each association was then defined as

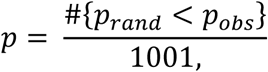

where #{*p*_rand_ < *p*_obs_} is the number of random p-values p_rand_ that are lower than the observed p-value p_obs_.

## 6 Author Contributions

S.Ta.: Conceptualisation, Methodology, Data Curation, Formal Analysis, Writing – original draft, Writing – review and editing; A.E.: Formal Analysis, Writing – review and editing; B.N.: Methodology, Data Curation, Writing – review and editing; L.S.: Data Curation, Writing – review and editing; M.O.H.: Methodology, Writing – review and editing; R.S., P.S., M.T., D.D., C.S., P.V.: Conceptualisation, Funding Acquisition, Writing – review and editing; D.P. and C.H.Y.: Conceptualisation, Methodology, Writing – review and editing; A.-L.P.: Conceptualisation, Methodology, Funding Acquisition, Writing – review and editing; All Authors: Writing – review and editing.

## 8 Acknowledgements

The authors thank the BIS participants for the generous contribution they have made to this project. The authors also thank current and past staff for their efforts in recruiting and maintaining the cohort and in obtaining and processing the data and biospecimens. We acknowledge Barwon Health, Murdoch Children’s Research Institute, and Deakin University for their support in the development of this research. The other members of the BIS Investigator Group are David Burgner, Lawrence Gray, Len Harrison, and Sarath Ranganathan. We thank John Carlin, Amy Loughman, Fiona Collier, Terry Dwyer, and Katie Allen for their past work as BIS investigators. We thank Andrea Gogos, Billi Newton, and Alicia Bjorksten for manuscript preparation.

