## Supplement for "Prenatal DEHP plastic chemical exposure increases the likelihood of child autism and ADHD symptoms through epigenetic programming"

##### This document includes:

1. **Supplementary Figure 1**: Barwon Infant Study (BIS) participant flowchart for ages 2 and 4 years
2. **Supplementary Table 1**: BIS cohort participant characteristics
3. Simulation analysis using synthetic data to demonstrate that the procedure for constructing MPSDEHP and Co-MN1 did not introduce bias towards significant mediation
    - **Supplementary Figure 2**
    - **Supplementary Figure 3**
4. **Supplementary Figure 4**: Expression levels by brain region of two key hormone receptors inferred *in silico* as upstream regulators of Co-$MN_1$ (*ESR1* and *NR3C1*)

### 1. Supplementary Figure 1

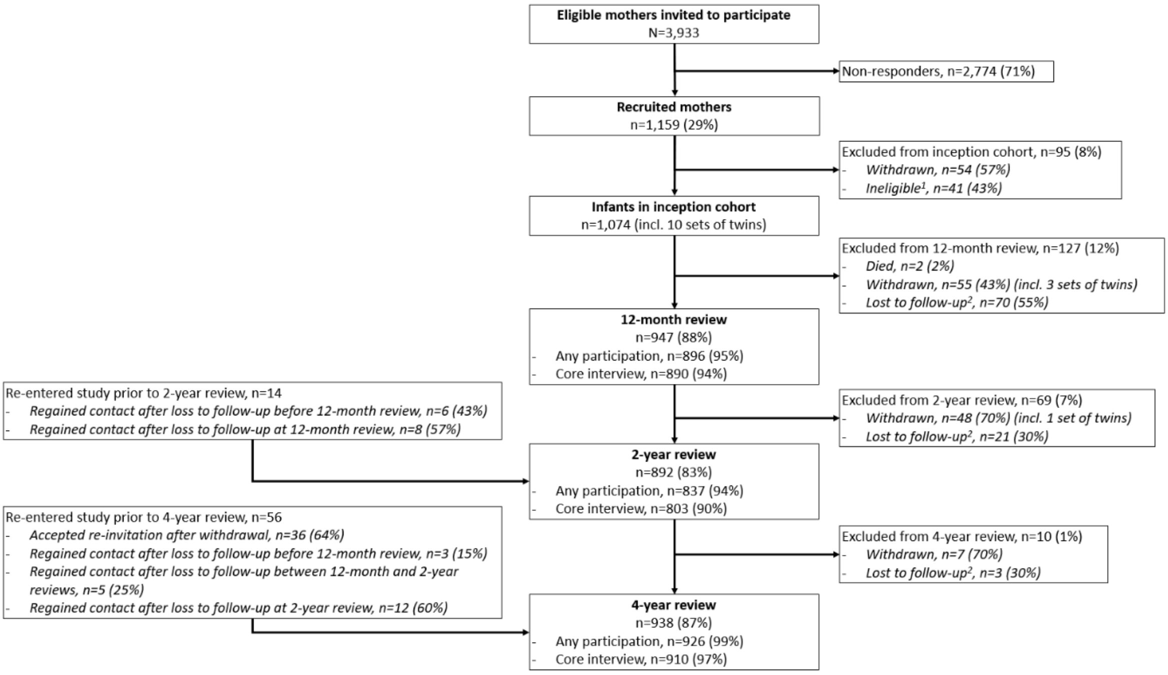


[1] Due to: no longer being resident in Barwon region (n=12), <32 weeks gestation (n=8), serious illness in first few days of life (n=7), major congenital disease (n=5), stillbirth (n=5), miscarriage (n=2), or cord blood stored privately (n=2)
[2] Loss to follow-up defined as missing two consecutive reviews

**Supplementary Fig. 1** | Barwon Infant Study (BIS) cohort participant flowchart up to age 4 years.

### 2. Supplementary Table 1

**Supplementary Table 1** | Key characteristics of participants in the Barwon Infant Study.

| | Cohort sample (*N* = 1074) | | Mediation sample (*N* ≤ 589) | |
|---|---|---|---|---|
| | ***N*** | **Mean (SD), median [IQR], % (n), geometric mean {GSD}** | ***N*** | **Mean (SD), median [IQR], % (n), geometric mean {GSD}** |
| **Parent and household factors** | | | | |
| Maternal age at conception (years) | 1074 | 31.33 (4.79) | 589 | 32.03 (4.26) |
| Paternal age at conception (years) | 1024 | 33.49 (5.85) | 563 | 33.88 (5.37) |
| SEIFA disadvantage (low tertile vs. other) | 1061 | | 583 | |
| Low (most disadvantage) | | 33.6 % (357) | | 29.2 % (170) |
| Medium | | 33.3 % (353) | | 32.8 % (191) |
| High (least disadvantage) | | 33.1 % (351) | | 38.1 % (222) |

| | | | | |
|---|---|---|---|---|
| Mean annual household income | 1041 | | 589 | |
| < AUD$25,000 | | 2.5 % (26) | | 0.7 % (4) |
| AUD$25,000–AUD$49,999 | | 9.5 % (99) | | 5.6 % (33) |
| AUD$50,000–AUD$74,000 | | 17.9 % (186) | | 18.2 % (107) |
| AUD$75,000–AUD$99,999 | | 25.6 % (266) | | 25.3 % (149) |
| AUD$100,000–AUD$149,999 | | 32.9 % (343) | | 36.7 % (216) |
| > AUD$150,000 | | 11.6 % (121) | | 13.6 % (80) |
| Maternal education | 1068 | | 589 | |
| Year 11 or less | | 8.6 % (92) | | 4.4 % (26) |
| Year 12 or equivalent | | 40.1 % (428) | | 37.0 % (218) |
| Bachelor or postgraduate degree | | 51.3 % (548) | | 58.6 % (345) |
| **Child genotyping** | | | | |
| Ethnicity principal component 1 | 1031 | -0.00 (0.03) | 589 | -0.00 (0.03) |
| **Prenatal factors** | | | | |
| Mother's pre-pregnancy BMI (kg/m²) | 927 | 25.38 (5.46) | 527 | 24.87 (5.00) |
| Maternal PSS score during pregnancy | 810 | 18.66 (6.99) | 455 | 17.85 (6.63) |
| Dietary energy (kJ/day) | 1016 | 7428.35 (2344.21) | 569 | 7464.08 (2209.81) |
| Walk days/week | 1009 | 3.98 (2.21) | 565 | 3.93 (2.13) |
| Maternal exposure to environmental tobacco smoke exposure in preconception or pregnancy (any vs. none) | 1049 | 16.9 % (177) | 578 | 14.4 % (83) |
| Mother smoked during pregnancy (any vs. none) | 1063 | 15.9 % (169) | 586 | 10.4 % (61) |
| Mother consumed alcohol during pregnancy (any vs. none) | 989 | 52.7 % (521) | 574 | 52.1 % (299) |
| Lone parent during pregnancy (yes vs. no) | 1071 | 4.0 % (43) | 589 | 2.7 % (16) |
| Multiparous (parity>1 vs. none) | 1073 | 55.3 % (593) | 589 | 56.9 % (335) |
| Gestational age at urine collection (weeks) | 847 | 36.27 (0.71) | 589 | 36.22 (0.64) |
| **Birth factors** | | | | |
| Gestational age at birth (weeks) | 1074 | 39.44 (1.52) | 589 | 39.64 (1.22) |
| Preterm birth (32-37 weeks vs rest) | 1074 | 4.4 % (47) | 589 | 1.4 % (8) |
| Multiple birth | 1074 | 0.9 % (10) | 589 | 0.0 % (0) |
| Mode of delivery | 1074 | | 589 | |

| | | | | |
|---|---|---|---|---|
| Unassisted vaginal birth | | 48.9 % (525) | | 49.4 % (291) |
| Assisted vaginal birth | | 19.9 % (214) | | 21.4 % (126) |
| Emergency caesarean section | | 16.7 % (179) | | 18.2 % (107) |
| Elective caesarean section | | 14.5 % (156) | | 11.0 % (65) |
| Maternal contamination of cord blood serum (any vs. none) | 936 | 4.6 % (43) | 589 | 0.0 % (0) |
| Resuscitation at birth (any vs. none) | 1074 | 0.8 % (9) | 589 | 0.8 % (5) |
| Apgar score at 1 min | 1059 | 9.00 [8.00, 9.00] | 581 | 9.00 [8.00, 9.00] |
| Birth weight (kg) | 1072 | 3.53 (0.52) | 589 | 3.59 (0.47) |
| Child sex (male vs. female) | 1074 | 51.7 % (555) | 589 | 50.8 % (299) |
| Glycoprotein acetyls, cord blood serum (mmol/l) | 909 | 0.73 {1.23} | 542 | 0.72 {1.19} |
| **Key prenatal exposures** | | | | |
| DEHP daily intake (µg/kg body weight/day) | 847 | 1.62 {2.08} | 589 | 1.63 {2.14} |
| DEHP daily intake > 3.26 (µg/kg body weight/day) | 847 | 1.8 % (15) | 589 | 2.0 % (12) |
| Maternal wholefoods diet | 1016 | 0.00 (1.00) | 569 | 0.11 (0.94) |
| **DNA methylation** | | | | |
| Co-$MN_1$ | 868 | 0.00 (0.03) | 589 | 0.00 (0.03) |
| $MPS_{DEHP}$ | 868 | -0.00 (1.00) | 589 | 0.02 (1.07) |
| **Neurodevelopmental assessments** | | | | |
| Child's age at CBCL/1.5–5 assessment (years) | 676 | 2.46 (0.15) | 489 | 2.46 (0.15) |
| CBCL/1.5-5 autism spectrum problems subscale (raw score) | 676 | 1.00 [0.00, 2.00] | 489 | 1.00 [0.00, 2.00] |
| CBCL 1.5-5 attention problems syndrome scale - externalising subscale (raw score) | 676 | 1.00 [0.00, 3.00] | 489 | 1.00 [0.00, 2.00] |
| CBCL/1.5-5 attention deficit hyperactivity problems subscale (raw score) | 676 | 3.00 [1.00, 5.00] | 489 | 3.00 [1.00, 5.00] |
| Child's age at SD assessment (years) | 791 | 4.16 (0.26) | 556 | 4.13 (0.23) |
| SDQ hyperactivity/inattention subscale | 791 | 3.00 [2.00, 5.00] | 556 | 3.00 [2.00, 5.00] |
| SDQ peer relationship problems subscale | 791 | 1.00 [0.00, 2.00] | 556 | 1.00 [0.00, 2.00] |

| | | | | |
|---|---|---|---|---|
| SDQ prosocial behaviour subscale | 792 | 8.00 [6.00, 9.00] | 556 | 8.00 [6.00, 9.00] |

NB. SD, standard deviation; IQR, interquartile range; GM, geometric mean; GSD, geometric standard deviation; SEIFA, Socio-Economic Indexes for Areas; DEHP, Di-(2-ethylhexyl) phthalate; CBCL, Child Behavior Checklist; SDQ, Strengths and Difficulties Questionnaire; ASD, autism spectrum disorder.

#### 3. Simulation analysis using synthetic data to demonstrate that the procedure for constructing $MPS_{DEHP}$ and $Co\text{-}MN_1$ did not introduce bias towards significant mediation

In this study, the putative mediators, $MPS_{DEHP}$ and $Co\text{-}MN_1$, were constructed from CpG sites filtered by association with the exposure, prenatal DEHP levels, making them closely related to this exposure. Given that prenatal DEHP exposure is associated with ASD and ADHD outcomes in the BIS cohort, an important consideration is whether this approach could create an artificial mediator-outcome association (simply because the mediator is a proxy for the exposure), potentially biasing the results towards significant mediated effects. To assess whether bias is introduced in mediation analysis when the mediator (*M*) serves as a proxy for the exposure (*E*), we conducted a simulation varying the correlation (ρ) between M and E. The parameter ρ (ranging from 0 to 1) controlled the extent to which *M* reflected *E*, where low values of ρ represented minimal correlation (i.e., *M* was independent of *E*) and high values represented *M* as an increasingly stronger proxy for *E*.

1. **Simulation Setup**: For each value of ρ, we conducted 50 simulations, each with a sample size of 100, generating *E* from a standard normal distribution and creating *M* as a linear combination of *E* and an independent noise term:

   $$M = \rho \times E + \sqrt{1-\rho^2} \times noise$$

   The outcome (*O*) was then simulated as a function of *E*, with additional noise, ensuring a significant exposure-outcome association to reflect the scenario in this study for prenatal DEHP exposure and the ASD and ADHD outcomes:

   $$O = 0.5 \times E + noise$$

2. **Mediation Analysis**: We used the `mediate` function from the `mediation` R package to estimate the Average Causal Mediation Effect (ACME) for each simulation, storing the p-values of the mediation effect for different values of ρ.

This procedure allowed us to examine whether increasing similarity between *M* and *E* (higher ρ) leads to biased significance in mediation results. Our findings from this simulation are summarised in Supplementary Figures 2 and 3 (below), which show that, contrary to concerns of bias toward significance, the likelihood of detecting significant mediation did not increase as M became a closer proxy for E.

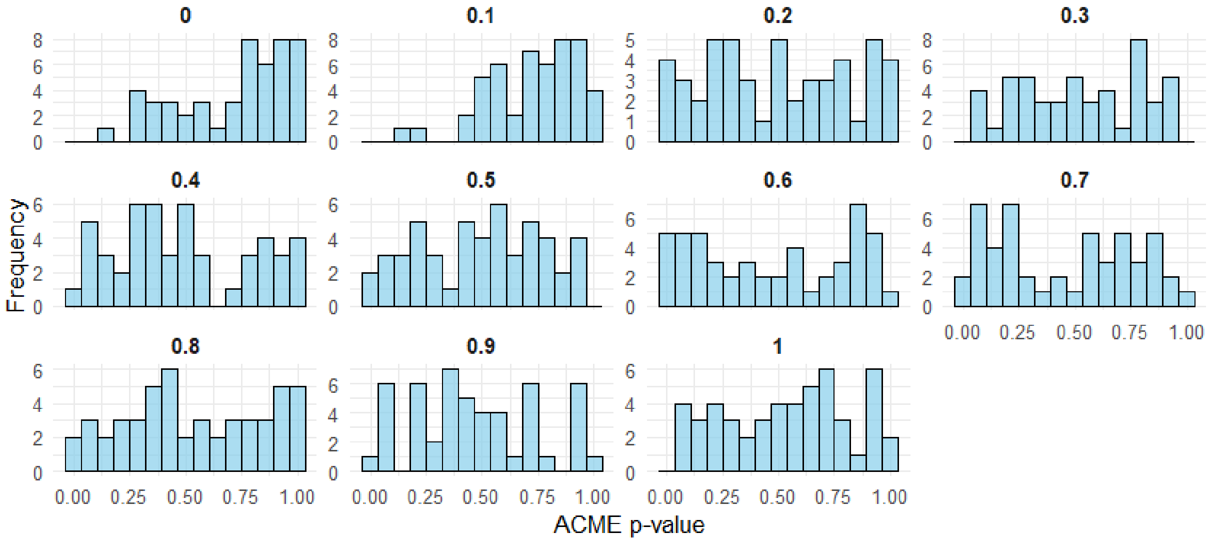


**Supplementary Fig. 2** | Distribution of estimated mediated effect (Average Causal Mediation Effect, or ACME) p-values across simulations for each value of *ρ* (from 0 to 1, in increments of 0.1). *ρ* represents the dependence of the mediator, M, on the exposure, E. Results show that p-values are uniformly distributed for high values of *ρ*, with no evidence of bias towards significance.

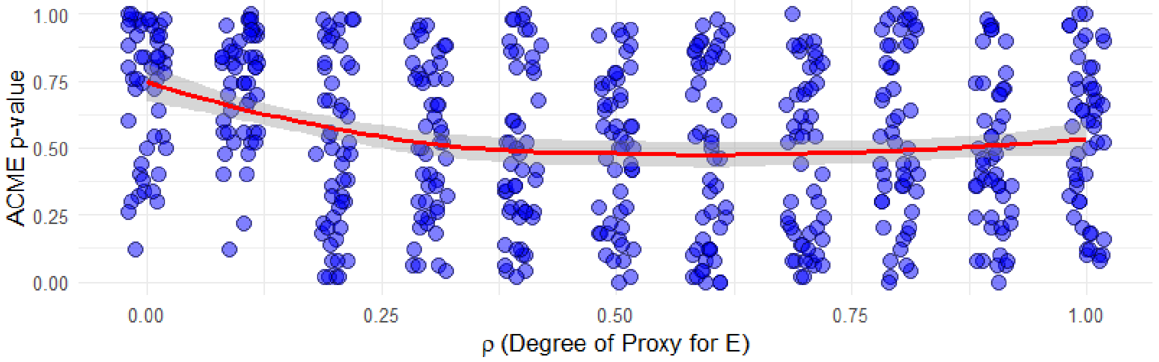


**Supplementary Fig. 3** | Distribution of mediated effect (Average Causal Mediation Effect, or ACME) p-values as *ρ* (reflecting the dependence of the mediator, *M,* on the exposure, *E*) increases. Mediated effect p-values are uniformly distributed around 0.5 for high values of *ρ*, demonstrating that no bias is introduced even when M is highly dependent on E. A shift away from 0.5 towards *non-significance* is evident for low values of ρ (0.0 to 0.3). This is because the exposure-mediator association, required to detect significant mediation, is generally missing here.

#### 4. Supplementary Figure 4

**a** *ESR1 expression*

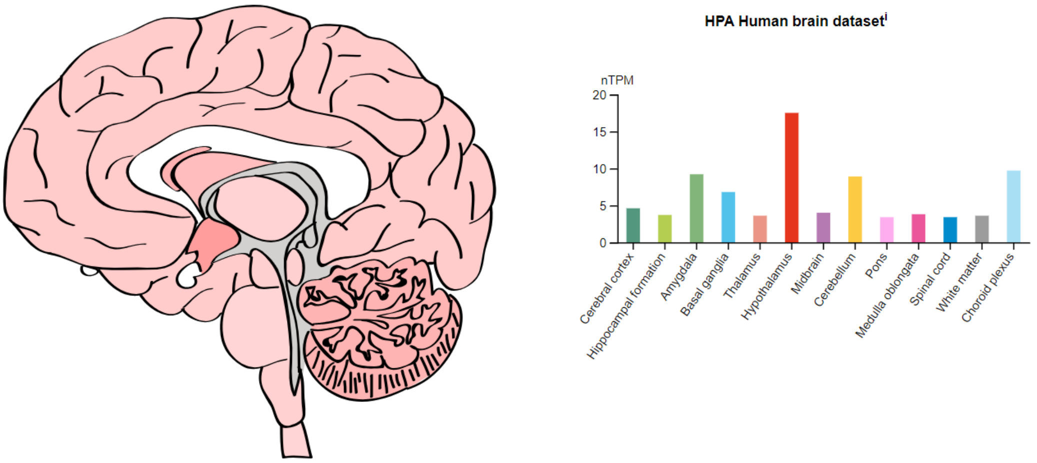


**b** *NR3C1 expression*

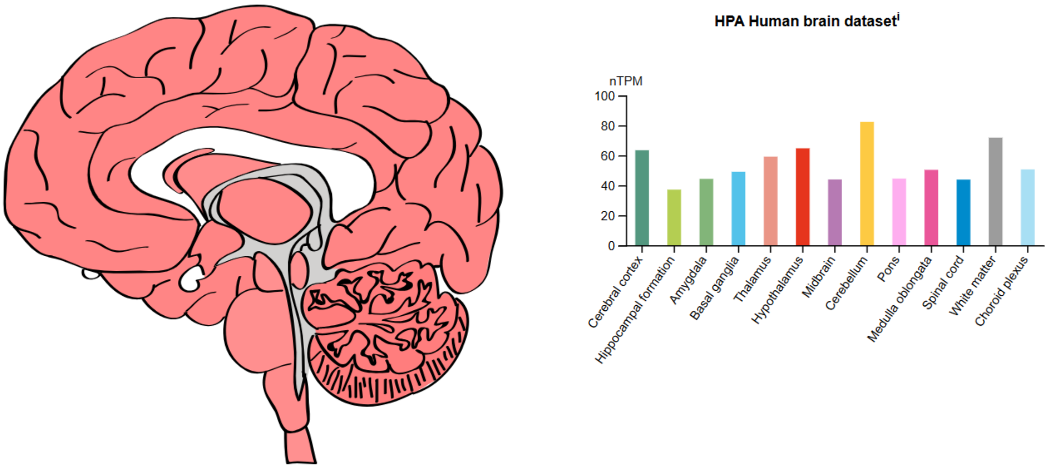


**Supplementary Fig. 4** | Expression levels by brain region of two key hormone receptors inferred *in silico* as upstream regulators of Co-MN$_1$. **a.** Expression of the Estrogen Receptor 1 (***ESR1***) gene. **b.** Expression of the Nuclear receptor subfamily 3 group C member 1 (***NR3C1***) gene, encoding the glucocorticoid receptor. The high expression of these genes in the cerebellum and in limbic-system structures may suggest elevated vulnerability in these regions to disruption of Co-MN$_1$. This is consistent with the independent cell-type enrichment analysis of Co-MN, which found an overrepresentation of marker genes in Co-MN$_1$ for these regions. Data shown here is from the Human Protein Atlas: https://www.proteinatlas.org/.
